# Genetic Surveillance Reveals Differential Evolutionary Dynamic of *Anopheles gambiae* Under Contrasting Insecticidal Tools used in Malaria control

**DOI:** 10.1101/2025.05.12.653619

**Authors:** Harun Njoroge, Lilian Namuli, Sanjay C Nagi, Anastasia Hernandez-Koutoucheva, Daniel P McDermott, Erin Knight, Samuel Gonahasa, Amy Lynd, Ambrose Oruni, Catherine Maiteki-Sebuguzi, Jimmy Opigo, Adoke Yeka, Agaba Katureebe, Mary Kyohere, Moses R Kamya, Grant Dorsey, Janet Hemingway, Sarah G Staedke, Chris Clarkson, Alistair Miles, Eric R. Lucas, Martin J Donnelly

## Abstract

Malaria, a febrile disease caused by the *Plasmodium* parasites and transmitted by mosquitoes, is a leading cause of mortality in children under 5 in endemic countries. The widespread deployment of insecticide-treated bed nets (ITNs) has significantly reduced malaria transmission, but rising levels of insecticide resistance threatens to halt the progress. Monitoring insecticide resistance is vital for effective vector control, particularly when deploying new tools. Understanding mosquito population responses to these interventions is crucial for guiding control programmes in making informed decisions about the selection, timing, and geographic deployment of tools. This genomic study investigates the demographic and evolutionary consequences on the malaria vector *Anopheles gambiae* of deploying standard ITNs (containing only pyrethroids) and pyrethroid-PBO nets (containing pyrethroids plus the synergist piperonyl butoxide) during a clinical trial in Uganda.

Despite substantial reductions in indoor mosquito densities in the clinical trial, estimates of nucleotide diversity (π) and linkage disequilibrium revealed no significant decline in effective population size, reflecting continued large population size even after effective control. Marked allele frequency shifts at resistance-associated loci indicated strong selection pressures driven by the interventions, with distinct selective dynamics between the two net types, highlighting alternative pyrethroid detoxification pathways in the presence of PBO. A duplication in the *Cyp9k1* gene significantly increased in frequency in populations exposed to pyrethroid-only nets but decreased in populations exposed to PBO-treated nets, suggesting that selection for over-expression of this gene is removed when this resistance mechanism is impacted by PBO. An alternative potential detoxification mechanism was selected within a region of the 2La chromosomal inversion on chromosome 2L, which encompasses the UDP-glucose 6-dehydrogenase gene. This variant consistently increased in frequency when exposed to PBO-treated nets. Additionally, pyrethroid-only nets selected for a novel locus on the X chromosome containing the diacylglycerol kinase gene, which is potentially linked to behavioural adaptations through its role in neurotransmission modulation.

Our findings underscore the importance of genomic surveillance in vector control, revealing distinct evolutionary dynamics of insecticide resistance mechanisms in the presence of PBO. While ITNs remain effective, the persistence and evolution of resistance-associated alleles highlight the need for adaptive and dynamic resistance management strategies. By integrating high-resolution genomic data with epidemiological and entomological monitoring, this study offers actionable insights to sustain malaria control efforts amid the ongoing challenge of insecticide resistance.

## Introduction

Malaria, a febrile disease caused by *Plasmodium* species and transmitted by *Anopheles* mosquitoes, accounts for over half a million deaths each year, predominantly in sub-Saharan Africa (WHO, 2024). Vector control methods using insecticides, particularly insecticide-treated bed nets (ITNs) and indoor residual spraying, are the main tools used in malaria control (Bhatt *et al*., 2015; Wilson *et al*., 2020). However, the emergence of insecticide resistance in *Anopheles* mosquitoes—the primary malaria vectors—necessitates innovative and sustainable strategies to achieve malaria elimination (Cotter *et al*., 2013; Mnzava *et al*., 2015). Effective malaria control requires rigorous evaluation of vector control tools through controlled trials and an understanding of how mosquito populations respond to them (Wilson *et al*., 2020).

Traditionally, malaria vector control has been monitored using entomological endpoints, including vector density, sporozoite infectivity rates, and entomological inoculation rates (EIR) (Morales-Pérezid *et al*., 2020; Traore *et al*., 2020). While these indicators remain the industry standards, they offer limited insights into the mechanisms underlying control effectiveness and are subject to biases with mosquito trapping techniques, environmental conditions, and technician proficiency, complicating comparisons between interventions (Rohani *et al*., 2016; Swai *et al*., 2023; Degefa, Yewhalaw and Yan, 2024).

Advances in genetic monitoring, informed by fields such as conservation genetics and pest management, provide opportunities to assess vector population dynamics, insecticide resistance, and intervention effectiveness with greater precision (Neafsey, Taylor and MacInnis, 2021; Schmidt, Endersby-Harshman and Hoffmann, 2021). Integrating genomic surveillance with traditional epidemiological indicators offers a promising approach to understanding vector population responses to control measures (Fouet, Atkinson and Kamdem, 2018). For instance, population genetics metrics such as F_ST_, heterozygosity, nucleotide diversity, and effective population size (N_e_) have been employed to assess the impact of ITNs, which when deployed can reduce vector abundance (Cartaxo, Ayres and Weetman, 2011; Athrey *et al*., 2012; O’Loughlin *et al*., 2016; Huang *et al*., 2023).

Genetic surveillance approaches can augment traditional phenotypic assays for insecticide resistance, which involve direct insecticide exposure, but can be influenced by environmental factors, mosquito physiology, and technician variability, limiting replicability (Praulins *et al*., 2024). Molecular diagnostics or targeted loci address some of these challenges, and while many markers are available for surveillance of insecticide resistance, they do not at present explain the full variation in resistance (Ranson *et al*., 2000; Jones *et al*., 2012; Weetman *et al*., 2018; Njoroge *et al*., 2022; Lucas *et al*., 2023) and are limited to discovering mutations at known loci.

In this study, we use whole-genome sequencing (WGS) of *An. gambiae s*.*s* collected during a large pragmatic cluster-randomized trial embedded within a national bed net distribution campaign in Uganda between 2017 to 2019, the LLINEUP trial (Staedke *et al*., 2020). The trial evaluated the effectiveness of standard pyrethroid only long-lasting insecticide treated bed nets (LLINs) against LLINs containing a pyrethroid + piperonyl butoxide (PBO) across 104 health sub-districts of Uganda. In Uganda, metabolic pyrethroid resistance mediated by cytochrome P450 enzymes is widespread and linked to a rapidly spreading resistance haplotype (Njoroge *et al*., 2022). Because the action of P450s is inhibited by PBO, the PBO + pyrethroid bed nets are expected to overcome the metabolic resistance, making them more effective. This was reflected in the epidemiological and entomological measurements which revealed a parasite prevalence ratio of 0.86 (0.74-1.00) and a vector density ratio of 0.38 (0.29-0.49) in the PBO + pyrethroid arm relative to pyrethroid only arm (Maiteki-Sebuguzi *et al*., 2023). Given the contrasting insecticidal pressure experienced by the mosquito population in the two arms, the trial offers an unprecedented opportunity to investigate the effects of large-scale vector interventions on the evolution and genetic makeup of a targeted vector population particularly to elucidate the evolutionary impact of PBO on cytochrome P450 mediated adaptive mechanism.

We hypothesized that (H_1_) the WGS data would reveal decreases in measures of vector effective population size across the entire study, (H_2_) reductions in N_e_ would be greater in the PBO arm compared to the pyrethroid-only arm, (H_3_) adaptive alleles would change in frequency across the study and (H_4_) distinct changes in frequencies of adaptive alleles would occur in the different arms, in particular for cytochrome P450 genes.

## Methods

### LLIN Evaluation in Uganda Project (LLINEUP): Study Design, Entomological Sampling, and Sample Processing

#### Study Design

The Long-Lasting Insecticidal Net Evaluation in Uganda Project (LLINEUP) was designed to assess the impact of LLINs, with and without PBO, on malaria indicators in Uganda. Conducted as a cluster-randomized trial embedded within a national LLIN distribution campaign (2017–2018), the study included two arms: one receiving standard pyrethroid nets and the other receiving PBO-pyrethroid impregnated nets.

A total of 104 health sub-districts (clusters) in Eastern and Western Uganda participated in the trial. Of these, 52 clusters were allocated PBO LLINs (PermaNet 3.0 and Olyset Plus), while 52 clusters received conventional LLINs (PermaNet 2.0 and Olyset Net). The primary outcome was malaria parasite prevalence in children aged 2–10 years, measured via microscopy. Secondary outcomes included anaemia prevalence, vector density, LLIN coverage, ownership, and usage. Details of the trial design, including sample size calculations and randomization methods, are available in previous publications (Staedke *et al*., 2019; Maiteki-Sebuguzi *et al*., 2023).

#### Entomological Sampling

Entomological surveillance was conducted to assess vector density, monitor insecticide resistance, and collect genomic data. Sampling was performed in all 104 clusters at baseline (prior to LLIN distribution), at 6-, 12- and 18 months but in 90 clusters at 25-months post-distribution. In each cluster, 10 households were randomly selected for indoor resting mosquito collection using Prokopack aspirators. Female *Anopheles* mosquitoes were identified morphologically, preserved in silica gel, and shipped to the Liverpool School of Tropical Medicine (LSTM) for further processing. Detailed sampling protocols are described elsewhere (Lynd *et al*., 2024).

#### Sample Processing

DNA was extracted from individual *Anopheles* mosquitoes using Nexttec extraction kits (Biotechnologie GmbH). Species identification was performed using a species-specific PCR assay targeting a SINE insertion site, distinguishing *An. gambiae s*.*s*., *An. arabiensis*, and *An. coluzzii* based on melt curve analysis (Chabi *et al*., 2019). This study focuses on *An. gambiae*, Uganda’s primary malaria vector at the time (Lynd *et al*., 2019). A subset of *An. gambiae* samples selected randomly within each entomological collection rounds and considering a good coverage across the health sub-districts were whole genome sequenced. Library preparation, sequencing, alignment, variant calling, and phasing were performed through the *An. Gambiae* 1000 genomes project pipeline (https://malariagen.github.io/vector-data/ag3/methods.html). Samples with coverage below 10x, males (determined by relative coverage on sex chromosome compared to autosomes), and specimens with evidence of contamination were excluded. A total of 1,013 mosquito whole genomes passed quality control. Due to limited samples from the 6, 12, and 18-month collection time points (PBO arm: N = 1, 14 and 1 respectively while Non PBO arm: N = 50, 154 and 126), analyses focused on comparisons between baseline (PBO arm: N = 185 and Non PBO arm: N = 207) and 25-month post-intervention populations (PBO arm: N = 53 and Non PBO arm: N = 222) (supplementary Table. 1).

#### Data Analysis

To evaluate the genomic impact of LLIN interventions on the population we structured the analysis to assess changes in population structure, size, and adaptive genomic responses, while explicitly addressing the four hypotheses. Population structure was assessed using principal component analysis (PCA) based on 100,000 randomly selected SNPs for each chromosome arm and genome wide windowed F_ST_ analysis to assess the degree of genetic differentiation within and between Eastern and Western Uganda populations.

Effective population size (N_e_) was estimated using linkage disequilibrium (LD) analysis (Nei, Maruyama and Chakraborty, 1975). LD values (r^2^) were calculated for 100,000 randomly sampled SNP pairs between chromosomes 2 and 3 (to exclude physically linked pairs) after excluding inversions regions (2Rb [2RL:17990000-32010000] and 2La [2RL:81535105-104555105]), singletons, non-segregating loci, and sites with missing data. Ne was calculated assuming a recombination rate of 0.5 (because SNP pairs were on separate chromosomes). Confidence intervals were calculated by 1000 iterations of sub-sampling 90% of the samples and recalculating Ne. To address H_1_, the LD and N_e_ of all pre-intervention samples were compared with values from size-matched post-intervention samples through random subsampling to account for LD’s sample size dependency and difference tested using Mann Whitney U test. For H_2_, populations exposed to PBO-treated and standard nets were analysed separately, with sample sizes equalized as with H_1_. A second method we used to assess changes in the population size was calculating temporal changes in nucleotide diversity (π), an indicator of recent demographic shifts, using the *malariagen-data* python package (Miles *et al*., 2024).

To test H_3_ and H_4_, we conducted genome-wide scans comparing baseline and post-intervention allele frequencies, restricting our analysis to high-confidence SNPs with good statistical power (biallelic SNPs with a minor allele frequency (MAF) >5% and no missing data, and passing Ag1000G’s site filters (https://malariagen.github.io/vector-data/ag3/methods.html#site-filtering). Generalized linear mixed models (GLMMs) were used to test the association between allele frequencies and intervention including intervention phase (baseline vs. 2 years post-intervention), geographic location (East vs. West) and net type (PBO nets vs. pyrethroid only nets) as fixed effects, and health sub-districts as a random effect (H_3_) (Brooks *et al*., 2017). Modelling was performed using the glmmTMB package (Brooks *et al*., 2017) in R (R Core Team, 2019) and P-values for intervention phase were calculated using the drop1 function, applying a Chi square test. Separate models for PBO and pyrethroid nets were fitted to test H_4_, and a Benjamini-Hochberg false discovery rate (FDR) correction was applied (Strimmer, 2008). Changes in frequency in SNPs with FDR-adjusted p-values <0.05 were considered significantly associated with intervention phase.

A complementary window-based approach using F_ST_ was applied to identify genomic regions undergoing differentiation between baseline and post-intervention populations. F_ST_ was calculated using the moving Patterson method in sliding windows of 1,000 SNPs with the Python package scikit-allel (https://zenodo.org/records/13772087). To identify significant outliers, we used a permutation-based approach like that described in Lucas et al., (2023). Briefly, the intervention phase values (pre-intervention, 2 years after intervention) were permuted across samples 1,000 times, and F_ST_ was recalculated for each permuted dataset. At each genomic window, empirical p-values were obtained by comparing the observed F_ST_ to the distribution of permuted values. To define significant peaks while minimizing false positives, we applied a thresholding approach based on the F_ST_ distribution. First, we determined the mode of the genome-wide F_ST_ distribution and calculated the difference between the mode and the minimum F_ST_ value. A peak threshold was set at three times this difference above the mode, ensuring that only the most extreme F_ST_ values were considered. Genomic windows were classified as significant if they exceeded this threshold and had an empirical p-value < 0.01.

To assess selective sweeps and adaptive responses, H12 statistics were calculated for phased haplotypes within 1,000-SNP sliding windows. ΔH12, defined as the difference in H12 values between post- and pre-intervention populations, was used to detect regions under positive selection. Positive ΔH12 values indicated reduced haplotype diversity post-intervention, suggesting increased selective pressure after the intervention, while negative values reflected increased haplotype diversity. To identify putative selective sweep regions, we defined peaks as windows with ΔH12 values exceeding three times the difference between the median and the 98th percentile of the genome-wide ΔH12 distribution. To assess statistical significance, we performed 1000 random permutations as described for F_ST_.

To identify and describe the haplotypes underlying significant signals of change over the intervention, we used hierarchical clustering of phased haplotypes using pairwise genetic distances (dxy). In each of the peaks where ΔH12 changed significantly, we used SNPs within genomic region encompassing 5% decay on both sides from the centre of H12 peak (*Supplementary Fig.1*). The centre of the peak was determined by fitting an exponential model on the calculated H12 to find the peak of the curve (https://malariagen.github.io/agam-selection-atlas/0.1-alpha3/index.html). Clusters of haplotypes forming a selective sweep were identified by cutting the tree using a threshold of dxy = 0.001. Each cluster was then considered as a haplotype allele. Associations between haplotype allele frequencies and intervention phases were assessed with GLMMs. We tested associations between these haplotype clusters and copy number variants (CNVs) calls at key metabolic resistance loci *Cyp6aa*-*Cyp6p*, and *Cyp9k1*(Miles *et al*., 2024). Because the CNV data are not phased, we identified tagging SNPs for the different CNV alleles at these loci through Spearman rank correlation of SNP calls and CNV calls at the sample level. For each CNV allele, the SNP with the highest correlation coefficient was used as a haplotype-level proxy for the CNVs.

#### Data and code availability

All samples were obtained from the LLINEUP trial which was approved by the Ugandan National Council for Science and Technology (UNCST Ref HS 2176) and the Liverpool School of Tropical Medicine (Ref 16-072). The epidemiological data from the LLINEUP trial can be accessed on ClinEpiDB ([LLINEUP Cluster Randomized Trial](https://clinepidb.org/ce/app/workspace/analyses/DS_7c4cd6bba9)). The variant data (SNPs and CNVs) used in this study are available through the MalariaGEN *Anopheles gambiae* 1000 Genomes Project (Ag1000G) and can be accessed using the malariagen-data Python package (https://malariagen.github.io/malariagen-data-python/v13.0.0/) or downloaded at https://malariagen.github.io/vector-data/ag3/download.html. The sample sets used are [“1288-VO-UG-DONNELLY-VMF00168”,”1288-VO-UG-DONNELLY-VMF00219”]. All code used for the analyses in this study is available in our public GitHub repository at https://github.com/vigg-lstm/LLINEUP-genomics. This repository contains scripts for data processing, population genetic analyses, and figure generation, ensuring full reproducibility of our results.

## Results

### Pre-Intervention Genetic Structure of *Anopheles gambiae* in Uganda

Principal component analysis (PCA) revealed no clear genetic differentiation between Eastern and Western Uganda populations (Supplementary Fig. 2). No structure was observed for any chromosome when excluding 2Rb and 2La chromosomal inversion regions. Inclusion of these regions revealed genetic structure by inversion genotypes, but not between East and West populations, confirming a high degree of genetic homogeneity across regions.

Nevertheless, weak isolation by distance may still exist that is not strong enough to be detected by PCA. To test this, we calculated F_ST_ between the geographical regions. We found localised significant differentiation in all the chromosomes particularly around genes linked to pyrethroid resistance suggesting a more rapid evolution within these loci than gene flow between the regions in Uganda (Supplementary Fig. 2). These results suggest that, while gene flow is extensive, regional differences exist at loci of recent selection. To account for this differentiation, location was incorporated as a fixed effect in subsequent generalized linear mixed models (GLMMs).

### H1 and H2 Genetic Analysis of Population Size Change After Bed Net Intervention

While the intervention led to a significant decline in indoor *An. gambiae* catch numbers in both arms of the LLINEUP trial in the first year and possible continued decline in the PBO arm (Supplementary table. 2), this decline was not detected in the genetic analysis of effective population size. LD is predicted to be larger in smaller populations due to more recent common ancestry (Hollenbeck, Portnoy and Gold, 2016), but we found no significant differences in LD or in the estimated N_e_ between baseline and post-intervention periods both with and without stratification by net type (Supplementary table. 3). Ne estimates derived from LD patterns were highly variable, with upper 95% confidence intervals extending to infinity and lower confidence intervals below 5,000, indicating the difficulty of reliable Ne estimation in very large populations, such as found in this species. Such high values were found both before and after intervention, indicating that despite the intervention, populations remain too large for accurate estimation of Ne. Similarly, nucleotide diversity showed no significant differences between baseline and post-intervention periods (Supplementary table. 3).

### H3 and H4 Genome-Wide Association Analysis Reveals Net Type- and Population-Specific Genetic Changes

Genome-wide SNP analysis identified 21 SNPs that exhibited significant frequency changes over the course of the intervention across the study sites (H3). Among these, a nonsynonymous SNP (2L:10334049, A>C, V390G) was detected in *AGAP005127*, which encodes RNA-binding protein 15. Seven SNPs were intronic, located within *CYP4G16* (X:22941281 and X:22941291), *AGAP029982* (3R:34361036), *C-type lectin* (X:18216512, X:18216520, X:18216526), and *Fatty acyl-CoA reductase* (X:23560372). The remaining SNPs were intergenic (Supplementary Fig. 3).

The intergenic SNPs were in the following regions on the genome; On chromosome 2R, five SNPs (2R:59036522 to 2R:59036642) were located 14 kb upstream of *Juvenile hormone diol kinase*, while two SNPs (2R:60709765 and 2R:60709812) were positioned 3.5 kb downstream of *AGAP004665* (a *CYP306A1*-like gene). Another SNP (2R:61318911) was identified 53 kb upstream of *AGAP004675*, a putative *muscarinic acetylcholine receptor 2*. Additional intergenic SNPs included two loci on chromosome 3R (3R:51526360 and 3R:51526384), located 3.7 kb upstream of *AGAP010243*, one SNP on 3L (3L:745991), positioned 24 kb upstream of *AGAP010324*, and another on 3L (3L:4240178), found 7.7 kb upstream of *AGAP010481*. On the X chromosome, SNP X:20244647 was located 6.5 kb upstream of *AGAP013064*.

In H4, we hypothesised that net-specific changes in SNP frequencies would be observed; however, no significant SNP frequency changes were detected when the trial arms were analysed separately. To complement single-SNP analyses, we utilised window-based methods (ΔH12) to detect broader signals of selection or differentiation. These methods revealed 5 key genomic regions where significant changes in allele frequencies occurred during the intervention (Fig.1 and supplementary Fig. 1): 2R:28463444– 28499726, 2L:2791320–2893275, 2L:34081017–34101131, X:9179019–9185374 and X:15216225– 15271654.

A net-specific change was clearly demonstrated in the X:15216225–15271654 region, which encompasses *Cyp9k1*, where ΔH12 showed a significant positive signal (Fig. 1) in both cohorts (Eastern and Western Uganda) that received standard bed nets, but the reverse was observed in both PBO bed net cohorts (Fig. 1). Haplotype clustering analysis linked the observed net-specific change to the main sweep haplotype allele in the X:15216225–15271654 genomic region, which significantly increased in frequency (FDR-corrected p = 0.04) in cohorts that received standard bed nets (Supplementary table. 4). This haplotype was associated with an increase in *Cyp9k1* copy number caused by the CNV allele *Cyp9k1*_Dup8 (Fig. 1). *Cyp9k1_*Dup8 increased in frequency in both cohorts that received standard bed nets (East 43% to 51% and West 84% to 93%) but decreased (East 56% to 54% and West 94% to 79%) in both PBO cohorts.

**Fig. 1.**
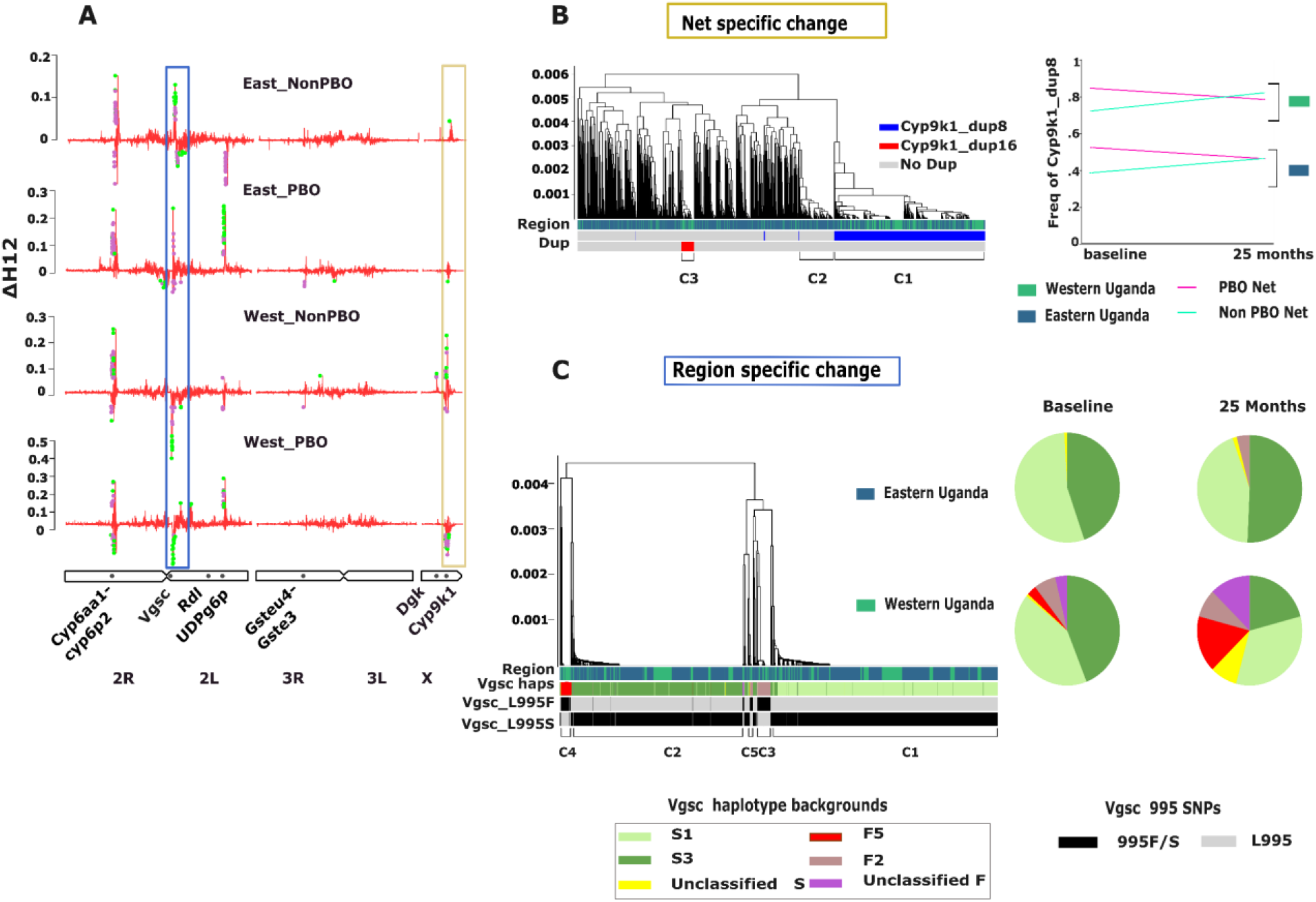
Targeted genomic responses to bed net interventions identified by genome-wide haplotype homozygosity (ΔH12) in Anopheles gambiae during the LLINEUP trial in Uganda. Genome-wide ΔH12 estimates (H12_end_– H12_baseline_) across four experimental cohorts from the LLINUP trial, plotted by chromosome arm. Positive ΔH12 values reflect reduced haplotype diversity post-intervention, suggesting directional selection. Coloured dots mark windows surpassing the permutation-based significance threshold (green: significant; purple: marginal). Labelled boxes denote regions of interest (blue: region specific change; Khaki: bed net specific change). **(B)** Dendrogram of haplotypes at the Cyp9k1 locus on chromosome X, highlighting net-specific selection. Distinct haplotypes associated with Cyp9k1 duplications (dup8, dup16) cluster with a lineage showing decreased frequency in PBO-treated cohorts and vice versa in Non PBO-treated cohorts (see ΔH12 peaks in panel A). The line plot (top right) tracks Cyp9k1 duplication frequency changes by region and treatment. **(C)** Dendrogram of the Vgsc locus on chromosome 2L, showing region-specific haplotype structure. Haplotype clusters correspond to known resistance alleles (Vgsc-L995F and Vgsc-L995S) and their haplotype backgrounds. Pie charts illustrate temporal shifts in Vgsc haplotype composition across regions and treatments, corresponding to ΔH12 dynamics in panel A.

Similarly, the X:9179019–9185374 locus also showed positive ΔH12 in standard bed net cohorts which was significant in Eastern Uganda, but no significant change in the PBO net cohorts from both locations in Uganda (Fig. 1 & Supplementary table. 4). The main haplotype in this region significantly increased in frequency in the overall population (H3: FDR-corrected p = 0.003) but this change was driven by the standard bed net cohorts (H4: FDR-corrected p = 0.0005) while in PBO bed net cohorts, the change was not significant (Supplementary table. 4). This genomic region lies within the intergenic region between *AGAP000516* (Enhancer of rudimentary protein) and *AGAP000519* (ATP dependent Diacylglycerol Kinase (*Dgk*)) (Fig.2). Neither gene has copy number variants, but *Dgk* has a missense mutation (L21F) which is associated with the swept haplotype (Fig. 2). The sweep is also associated with other non-coding SNPs from the genomic region (Supplementary table 5).

**Fig. 2.**
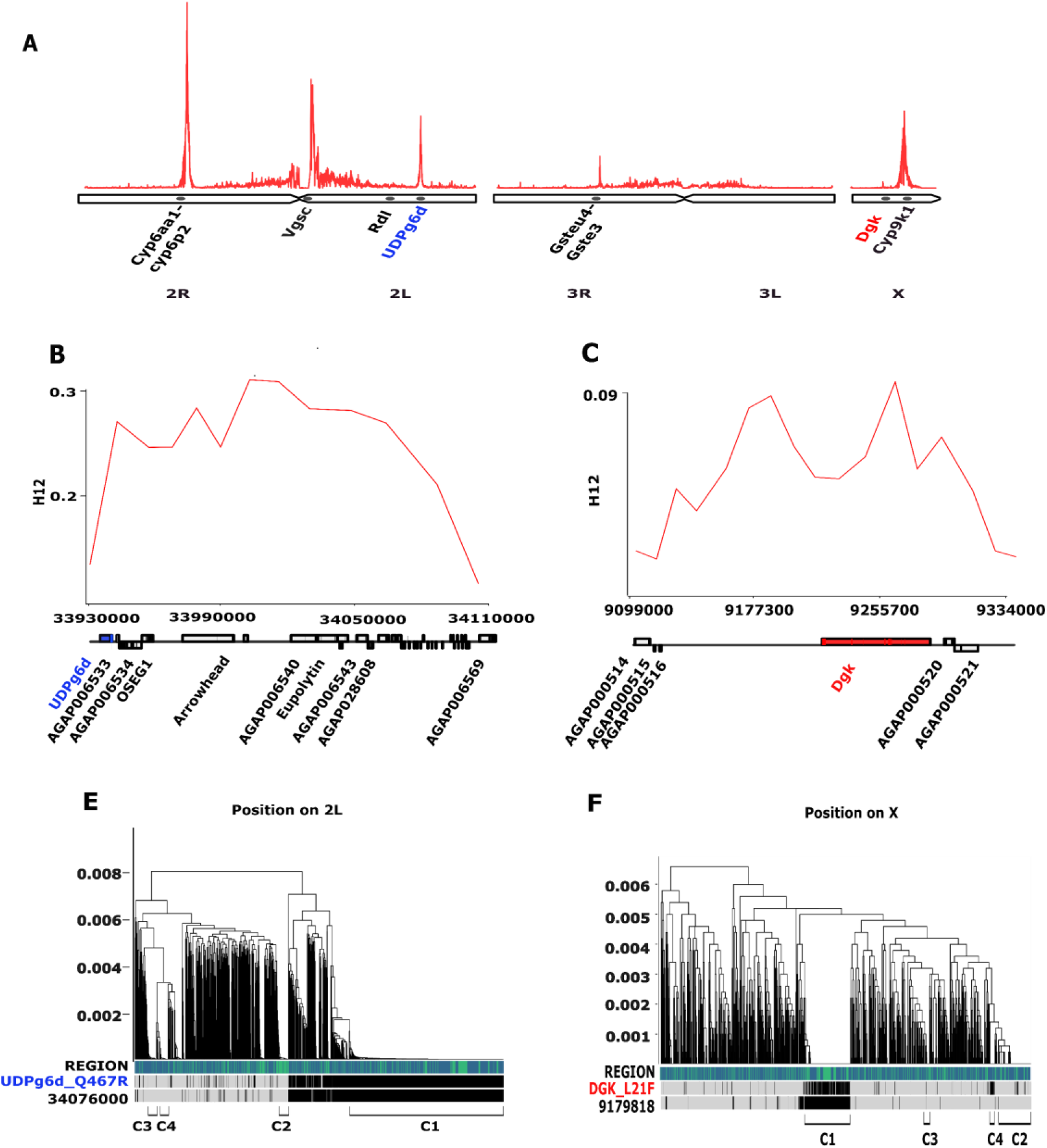
Novel selective sweeps impacted by bed net interventions reveal putative drivers of resistance evolution in Anopheles gambiae. (A) Genome-wide H12 scan of Ugandan An. gambiae populations identifies regions under recent selection. In addition to known insecticide resistance loci, a prominent novel sweep on chromosome 2L (∼34 Mb) includes the metabolic gene UDP-glucose 6-dehydrogenase (UDPg6p), and a sweep on the X chromosome (∼9.2 Mb) spans the diacylglycerol kinase (Dgk) gene. **(B–C)** Local H12 profiles for the 2L (B) and X (C) regions show elevated haplotype homozygosity across windows surrounding the candidate genes. **(D–F)** Dendrograms of haplotypes in the same 2L (E) and X (F) sweep regions highlight the dominant haplotype clusters associated with the selective sweeps. Candidate genes and putative tagging SNPs are annotated below each panel.

A change in frequency in a swept haplotype within the 2La chromosomal inversion on chromosome 2L (2L:34081017–34101131) also followed a net-specific trend, but opposite to that observed in the two regions on the X chromosome (Fig. 1). A significant increase in the main haplotype allele (FDR-adjusted p = 0.003) occurred in cohorts that received PBO bed nets, whereas no significant change was detected in the standard bed net cohorts. The main haplotype is not linked to copy number variations which are absent in the genes within the region however a missense SNP in 2L:33945793 *(AGAP006532*_Q467R) in *UDP glucose 6 dehydrogenase* (Fig.2) and other SNPs were associated with the haplotype (Supplementary Table. 5).

The 2R:28463444–28499726 and 2L:2791320–2893275 swept haplotype changes were not net-specific, aligning with the H3 hypothesis (Fig. 1). In both bed net cohorts, the swept haplotypes in the 2R:28463444– 28499726 genomic region where the *Cyp6aa1-Cyp6p2* gene cluster is located, increased in frequency (Fig. 1). Recently, a major haplotype linked to a duplication in the *Cyp6aa1* gene (Supplementary Fig. 4), a SNP (I236M) in the *Cyp6p4* gene and an insertion of a transposable element was identified in the same genomic region. The haplotype is highly associated in pyrethroid resistance in East and Central Africa (Njoroge *et al*., 2022). Despite having significant positive ΔH12 (Fig. 1), the main haplotype did not significantly increase in frequency in both bed net cohorts during the intervention but the second most common haplotype which lacks duplication of the *Cyp6aa1* gene (Supplementary Fig. 4) decreased significantly (FDR adjusted p = 0.02) in frequency (Supplementary table. 4) implying it was less advantageous in the presence of the interventions.

In the 2L:2791320–2893275 region around the *Vgsc* gene, the main haplotypes (S1 and S3 (Clarkson *et al*., 2021)) which are associated with the *Vgsc-*995S mutation (Fig. 1) increased in frequency in both PBO and standard bed net cohorts in Eastern Uganda, while the opposite trend occurred in Western Uganda, where a minor haplotype (F5) linked to *Vgsc*-995F (Fig. 1) increased in frequency as the main (*Vgsc-*995S) haplotypes declined (Supplementary table 4 and 6). This *Vgsc*-995F haplotype was confined to Western Uganda (Fig. 1) where it increased in frequency from 2% to 8.3% in cohorts that received standard bed nets while in PBO cohorts it increased from 2.7% to 14.3%.

## Discussion

This study provides a genomic insight into the adaptive responses of *An. gambiae* populations to large-scale bed net interventions, highlighting the selective pressures exerted by insecticide-based vector control. Using genome-wide SNP analyses, haplotype clustering, and diversity metrics, we provide evidence of population-specific and intervention-specific genetic changes, particularly at loci associated with pyrethroid resistance.

### Genetic Diversity and Population Structure

Contrary to our initial expectations based on the entomological outcomes of the LLINEUP trial (Staedke *et al*., 2020; Maiteki-Sebuguzi *et al*., 2023), nucleotide diversity (π), linkage disequilibrium (LD), and effective population size (N_e_) estimates (H_1_ and H_2_) did not reveal significant reductions in effective population size following the intervention. This contrasts with the reported reductions in indoor *Anopheles* densities across all cohorts, with greater reductions in the PBO net cohorts. While the number of captured mosquitoes declined, our results indicate that the population remains large enough that genetic signatures of a reduced population size could not be detected.

The LD method is most effective at detecting changes in Ne when population size is small, becomes increasingly inaccurate in large populations, such as those with high genetic diversity and extensive breeding populations characteristic of *An. gambiae* (Leffler *et al*., 2012; The Anopheles gambiae 1000 Genomes Consortium, 2017). In such cases, the method often fails to provide meaningful estimates, frequently reporting infinite N_e_ values. Similar limitations have been observed in other studies estimating N_e_ in *Aedes aegypti, Culex quinquefasciatus*, and the Miami blue butterfly where LD-based estimates often yielded unrealistic values compared to temporal and coalescent-based approaches (Saarinen, Austin and Daniels, 2010; Saarman *et al*., 2017; Huang *et al*., 2023).

While comparing LD before and after an intervention without explicitly estimating Ne can provide insight into population size changes, our data did not indicate any significant reduction during the trial (Supplementary Table 3), again indicating that the population remained too large for fluctuations to result changes in LD. In addition, diversity statistics remain unchanged during the intervention thus the data supports the effective population size remained large throughout the trial without any detectable changes. The steady population despite the deployment of control tools can result from high gene flow, as evidenced by PCA and differentiation analyses in our study, suggesting that minimal barriers to inter-regional migration may have allowed for the replenishment of localized diversity losses. Alternatively, the decline in trapped mosquitoes resting indoors as reported in the LLINEUP trial (Maiteki-Sebuguzi *et al*., 2023) may also result from behavioural avoidance driven by interventions, including shifts in feeding or resting preferences, which have been observed in other field studies (Prussing *et al*., 2018; Degefa *et al*., 2021; Machani *et al*., 2022; Omondi *et al*., 2023).

### Locus-Specific Selection Pressures

Although genetic diversity and effective population size apparently remained constant, our results show that the population was nonetheless impacted by interventions, with selection acting on specific loci or haplotypes. Significant changes in allele frequencies at resistance-associated loci highlight the intense selective pressures imposed by bed nets. Regions encompassing genes known to be associated with resistance, such as the pyrethroid target site *Vgsc* (Grigoraki *et al*., 2021), and metabolic resistance gene regions *Cyp6aa1-Cyp6p2*, and *Cyp9k1* (Vontas *et al*., 2018; Njoroge *et al*., 2022) exhibited patterns consistent with positive selection, aligning with their established roles in target-site and metabolic resistance mechanisms.

For instance, the increased frequency of the main haplotype in the *Cyp6aa1-Cyp6p2* region, which co-occurs with *Cyp6aa1* duplication and *Cyp6p4*-*236M* mutations, underscores its adaptive significance. This haplotype, previously identified in East and Central Africa, confers enhanced metabolic detoxification capacity, facilitating survival in the presence of pyrethroids (Njoroge *et al*., 2022). Similarly, the dynamics at the *Vgsc* locus reveal the persistence of knockdown resistance alleles. Though our sample size was modest (Supplementary Table 1), the observed shifts in *Vgsc* haplotypes corroborate a model showing the flow of the *Vgsc-995F* from the west into Western Uganda (Lynd *et al*., 2024). We used the same tagging SNPs and haplotype allocation method as Lucas et al., (2019) to identify the actual knockdown resistance haplotypes that were changing in frequency during the intervention. These origins include S1 to S5 which are the different haplotype backgrounds of the *Vgsc*-995S mutation and F1 to F5 the haplotype backgrounds of *Vgsc*-995F. In East Africa, the main haplotypes are S1 and S3. The haplotype background analysis revealed the F5 haplotype which was originally identified in Cameroon, Gabon and Democratic Republic of Congo (Lucas *et al*., 2019; Clarkson *et al*., 2021) increased in frequency during the intervention in Western Uganda while S1 and S3 haplotypes decreased in frequency (Fig. 1). The replacement of *Vgsc-995S* haplotype backgrounds with *Vgsc-995F* haplotype reflects the adaptive advantage of the latter (Lynd *et al*., 2010), consistent with its near fixation of *Vgsc-995F* in West African populations highly resistant to pyrethroids (Kouamé *et al*., 2023).

### Net-Specific Selective Pressures

We observed significant net-specific changes in the frequencies of major alleles within three genomic regions: *2La-34Mb* (2L:34,081,017–34,101,131), *AGAP000516-Dgk* (X:9,179,019–9,185,374), and *Cyp9k1* (X:15,216,225–15,271,654). Notably, a significant increase in the frequency of the primary haplotype in the *2La-34Mb* region occurred within the intervention arm that received PBO + pyrethroid bed nets. This suggests that PBO nets impose distinct selective pressures on this genomic region. While the selective sweep at *2La-34Mb* has previously been found, the selection pressure driving the sweep is unknown (The Anopheles gambiae 1000 Genomes Consortium, 2017), the increased frequency of this haplotype during the intervention indicates that adaptive genetic variants in this region may offer alternative resistance mechanisms when cytochrome P450-mediated resistance is inhibited.

The 2La chromosomal inversion is primarily associated with thermal and desiccation tolerance (Gray *et al*., 2009; Rocca *et al*., 2009). It has also been linked to dieldrin resistance, which is related to the segregation of the 296G/S mutations in the *GABA-gated chloride channel* gene (*GABA*) located within the inversion (Adeogun *et al*., 2019; Grau-Bové *et al*., 2020). However, the *GABA* gene (2L:25,363,652–25,434,556) lies outside the swept region identified in this study, and the 296G/S mutations are absent in our population, making it unlikely that this gene contributed to the observed positive selection.

Mosquitoes carrying the standard form of the 2La inversion possess thicker cuticles, which not only reduce desiccation (Reidenbach *et al*., 2014), but can also act as a barrier to insecticide uptake (Balabanidou *et al*., 2016). This observation aligns with findings from Kenya, where the frequency of the standard chromosomal form increased from 0% to 76% between 1994 and 2011, coinciding with the rise in pyrethroid bed-net coverage from 0% to over 90%. This change occurred independently of climatic factors (Matoke-Muhia *et al*., 2016). Similarly, in our Ugandan population, the swept haplotype was found exclusively in mosquitoes carrying the standard 2La chromosome. This strong association makes it challenging to discern whether the increase in frequency was driven by the swept haplotype itself or the whole standard 2La chromosome form.

Among the genes within the swept region, none are directly linked to cuticle metabolism, making it unlikely that the sweep is associated with cuticle thickening. However, *UDP-glucose 6-dehydrogenase*, a gene involved in detoxification, is located within this region. This gene encodes an enzyme that catalyses the oxidation of UDP-glucose to UDP-glucuronic acid, which is subsequently conjugated by UDP-glucosyltransferases with endogenous or exogenous compounds, facilitating their deactivation or excretion (Real, Ferre and Chapa, 1991; Hung *et al*., 2007). Logan et al., (2024) demonstrated that this pathway is involved in pyrethroid detoxification in *An. gambiae* and *An. funestus*. Thus, this pathway may be an important alternative when cytochrome P450 enzymes are inhibited by PBO, providing an explanation for the increased frequency of this haplotype.

In contrast, standard pyrethroid-only nets maintained selective pressures favouring metabolic resistance, as evidenced by changes in the *Cyp9k1* region. *Cyp9k1*, a cytochrome P450 gene, is a key pyrethroid metabolizer, explaining the increased frequency of this haplotype during the intervention (Vontas *et al*., 2018). This gene is duplicated in Uganda where the main CNV allele is *Cyp9k1*_Dup8 with a secondary minor allele, *Cyp9k1_*Dup16. We found that *Cyp9k1*_ Dup8 is fully linked to the main haplotype that was increasing in frequency during the bed net intervention (Fig. 1). The elevated copy number in *Cyp9k1* likely results in additional enzyme copies, enhancing the overall capacity for pyrethroid metabolism and thus increasing resistance. However, in the presence of PBO, which inhibits cytochrome P450 enzymes (Jones, 1998), additional copies of *Cyp9k1* may offer no advantage. This could explain the observed negative selection of the haplotype in populations exposed to PBO + pyrethroid bed nets (Fig. 1). Our findings suggest that PBO may disable *Cyp9k1* detoxification of pyrethroids sufficiently. However, alternative mechanisms, such as the proposed involvement of UDP-glucosyltransferases, may be recruited to compensate for this loss.

A novel swept locus X:9,179,019–9,185,374 on the X chromosome also exhibited significant increase in frequency over the intervention in the pyrethroid-only cohort. Unlike the *Cyp9k1* region, no opposite change was detected in the PBO + pyrethroid cohorts. This locus flanked by several genes: *Agap000513* (*dipeptidase E), Agap000515, Agap000516* (*Enhancer of rudimentary protein*), and *Agap000519* (*diacylglycerol kinase, Dgk*), none of which have been definitively shown to play a direct role in adaptation to pyrethroids. However, there are reasons to believe that *Dgk* may play a role in resistance. It is involved in modulating levels of secondary lipid messengers, such as diacylglycerol (DAG) and phosphatidic acid (PA), which influence key biological processes, including neurotransmission, lipid metabolism, protein kinase C activity, vesicle trafficking, and the regulation of membrane protein function (Topham, 2006). The roles of DGK in neurotransmission and vesicle trafficking may indirectly contribute to pyrethroid resistance, but this remains speculative.

### Implications for Resistance Management and Future Directions

Our findings emphasize the interplay between intervention type and mosquito genetic responses. The persistence of resistance-associated alleles, even with novel nets containing a resistance breaking synergist, underscores the challenge of overcoming pre-existing adaptations. Additionally, the identification of loci like *Dgk* and SNPs within intergenic or introns, which lack established links to resistance, raises questions about alternative adaptive mechanisms, including behavioural resistance or regulatory element involvement. These loci warrant further investigation to understand their roles in the adaptive landscape of *An. gambiae*.

While this study provides a comprehensive analysis of genetic responses to bed net interventions, certain limitations should be noted. Intermediate changes at 6-, 12-, and 18-months post-intervention could not be analysed due to limited sample sizes. This is the period when biggest drops in catch numbers were seen and likely the highest chance of finding a drop in Ne, but they could not be analysed because of small sample size. Key changes may have been missed due to gene flow from non-intervention regions or reduced selection pressure as insecticide efficacy waned. Expanding the temporal and spatial scope of sampling is essential for a comprehensive understanding of resistance dynamics.

## Conclusions

Genomic surveillance should remain central to malaria control programs, enabling the early detection of emerging resistance alleles and informing adaptive strategies (Weetman and Donnelly, 2015). In regions like Uganda, where *An. gambiae* populations exhibit both high genetic homogeneity and region-specific selective pressures, targeted interventions accounting for these dynamics could sustain control measures’ efficacy. By integrating genomic surveillance with epidemiological and entomological monitoring, we can develop more sustainable, evidence-based strategies to combat malaria and insecticide resistance. Our findings underscore the advantages of whole-genome sequencing in identifying adaptive responses missed by approaches that use only known resistance markers (Lynd *et al*., 2024) emphasising the importance of being integrated in control programs. Tracking changes in population structure, diversity, and resistance allele frequencies provides early warning signals of control failure, guiding the strategic deployment of resistance management tactics in an era of rising resistance (Neafsey, Taylor and MacInnis, 2021).

## Notes

### Competing Interest Statement

The authors have declared no competing interest.

### Funding

- National Institute of Allergy and Infectious Diseases of the National Institutes of Health: R01AI116811
- Bill & Melinda Gates Foundation: OPP1210750
- MRC DTP studentship: MR/R015678/1

## References

Adeogun, A.O. et al. (2019) Test for association between dieldrin resistance and 2La inversion polymorphism in Anopheles coluzzii from Lagos, Nigeria, Tropical Biomedicine.

Athrey, G. et al. (2012) ‘The Effective Population Size of Malaria Mosquitoes: Large Impact of Vector Control’, PLOS Genetics, 8(12), pp. e1003097.. Available at: 10.1371/journal.pgen.1003097.

Balabanidou, V. et al. (2016) ‘Cytochrome P450 associated with insecticide resistance catalyzes cuticular hydrocarbon production in Anopheles gambiae’, Proc. Natl. Acad. Sci., 113(33), p. 201608295. Available at: 10.1073/pnas.1608295113.

Bhatt, S. et al. (2015) ‘The effect of malaria control on Plasmodium falciparum in Africa between 2000 and 2015.’, Nature, 526(7572), pp. 207–11. Available at: 10.1038/nature15535.

Brooks, M.E. et al. (2017) ‘glmmTMB balances speed and flexibility among packages for zero-inflated generalized linear mixed modeling’, The R journal, 9(2), pp. 378–400.

Cartaxo, M.F.S., Ayres, C.F.J. and Weetman, D. (2011) ‘Loss of genetic diversity in Culex quinquefasciatus targeted by a lymphatic filariasis vector control program in Recife, Brazil’, Transactions of the Royal Society of Tropical Medicine and Hygiene, 105(9), pp. 491–499. Available at: 10.1016/j.trstmh.2011.05.004.

Chabi, J. et al. (2019) ‘Rapid high throughput SYBR green assay for identifying the malaria vectors Anopheles arabiensis, Anopheles coluzzii and Anopheles gambiae s.s. Giles’, PLoS ONE, 14(4). Available at: 10.1371/journal.pone.0215669.

Clarkson, C.S. et al. (2021) ‘The genetic architecture of target-site resistance to pyrethroid insecticides in the African malaria vectors Anopheles gambiae and Anopheles coluzzii’, Molecular ecology, 30(21), pp. 5303–5317.

Cotter, C. et al. (2013) ‘The changing epidemiology of malaria elimination: new strategies for new challenges.’, Lancet, 382(9895), pp. 900–11. Available at: 10.1016/S0140-6736(13)60310-4.

Degefa, T. et al. (2021) ‘Patterns of human exposure to early evening and outdoor biting mosquitoes and residual malaria transmission in Ethiopia’, Acta Tropica, 216. Available at: 10.1016/j.actatropica.2021.105837.

Degefa, T., Yewhalaw, D. and Yan, G. (2024) ‘Methods of sampling malaria vectors and their reliability in estimating entomological indices in Africa’, Journal of Medical Entomology, p. tjae015. Available at: 10.1093/jme/tjae015.

Fouet, C., Atkinson, P. and Kamdem, C. (2018) ‘Human Interventions: Driving Forces of Mosquito Evolution’, Trends in Parasitology. Elsevier Ltd, pp. 127–139. Available at: 10.1016/j.pt.2017.10.012.

Grau-Bové, X. et al. (2020) ‘Evolution of the insecticide target Rdl in african anopheles is driven by interspecific and interkaryotypic introgression’, Molecular Biology and Evolution, 37(10), pp. 2900–2917. Available at: 10.1093/molbev/msaa128.

Gray, E.M. et al. (2009) ‘Inversion 2La is associated with enhanced desiccation resistance in Anopheles gambiae’, Malaria Journal, 8(1). Available at: 10.1186/1475-2875-8-215.

Grigoraki, L. et al. (2021) ‘CRISPR/Cas9 modified An. Gambiae carrying kdr mutation L1014F functionally validate its contribution in insecticide resistance and combined effect with metabolic enzymes’, PLoS Genetics, 17(7). Available at: 10.1371/journal.pgen.1009556.

Hollenbeck, C.M., Portnoy, D.S. and Gold, J.R. (2016) ‘A method for detecting recent changes in contemporary effective population size from linkage disequilibrium at linked and unlinked loci’, Heredity, 117(4), pp. 207–216. Available at: 10.1038/hdy.2016.30.

Huang, X. et al. (2023) ‘Effective population size of Culex quinquefasciatus under insecticide-based vector management and following Hurricane Harvey in Harris County, Texas’, Frontiers in Genetics, 14. Available at: 10.3389/fgene.2023.1297271.

Hung, R.J. et al. (2007) ‘Comparative analysis of two UDP-glucose dehydrogenases in Pseudomonas aeruginosa PAO1’, Journal of Biological Chemistry, 282(24), pp. 17738–17748. Available at: 10.1074/jbc.M701824200.

Jones, C.M. et al. (2012) ‘Footprints of positive selection associated with a mutation (N1575Y) in the voltage-gated sodium channel of Anopheles gambiae’, PNAS, 109(17). Available at: 10.1073/pnas.1201475109.

Jones, D.Glynne. (1998) Piperonyl butoxide : the insecticide synergist. Academic Press.

Kouamé, R.M.A. et al. (2023) ‘Widespread occurrence of copy number variants and fixation of pyrethroid target site resistance in Anopheles gambiae (s.l.) from southern Côte d’Ivoire’, Current Research in Parasitology and Vector-Borne Diseases, 3. Available at: 10.1016/j.crpvbd.2023.100117.

Leffler, E.M. et al. (2012) ‘Revisiting an Old Riddle : What Determines Genetic Diversity Levels within Species ?’, PLoS Biol., 10(9). Available at: 10.1371/journal.pbio.1001388.

Logan, R.A.E. et al. (2024) ‘Uridine diphosphate (UDP)-glycosyltransferases (UGTs) are associated with insecticide resistance in the major malaria vectors Anopheles gambiae s.l. and Anopheles funestus’, Scientific Reports, 14(1). Available at: 10.1038/s41598-024-70713-y.

Lucas, E.R. et al. (2019) ‘A high throughput multi-locus insecticide resistance marker panel for tracking resistance emergence and spread in Anopheles gambiae’, Scientific reports, 9(1), pp. 1–10.

Lucas, E.R. et al. (2023) ‘Genome-wide association studies reveal novel loci associated with pyrethroid and organophosphate resistance in Anopheles gambiae and Anopheles coluzzii’, Nature Communications, 14(1). Available at: 10.1038/s41467-023-40693-0.

Lynd, A. et al. (2010) ‘Field, genetic, and modeling approaches show strong positive selection acting upon an insecticide resistance mutation in anopheles gambiae s.s.’, Molecular Biology and Evolution, 27(5), pp. 1117–1125. Available at: 10.1093/molbev/msq002.

Lynd, A. et al. (2019) ‘LLIN Evaluation in Uganda Project (LLINEUP): A cross-sectional survey of species diversity and insecticide resistance in 48 districts of Uganda’, Parasites and Vectors. BioMed Central Ltd. Available at: 10.1186/s13071-019-3353-7.

Lynd, A. et al. (2024) ‘LLIN Evaluation in Uganda Project (LLINEUP)–effects of a vector control trial on Plasmodium infection prevalence and genotypic markers of insecticide resistance in Anopheles vectors from 48 districts of Uganda’, Scientific Reports, 14(1). Available at: 10.1038/s41598-024-65050-z.

Machani, M.G. et al. (2022) ‘Behavioral responses of pyrethroid resistant and susceptible Anopheles gambiae mosquitoes to insecticide treated bed net’, PLoS ONE, 17(4 April). Available at: 10.1371/journal.pone.0266420.

Maiteki-Sebuguzi, C. et al. (2023) ‘Effect of long-lasting insecticidal nets with and without piperonyl butoxide on malaria indicators in Uganda (LLINEUP): final results of a cluster-randomised trial embedded in a national distribution campaign’, The Lancet Infectious Diseases, 23(2), pp. 247–258. Available at: 10.1016/S1473-3099(22)00469-8.

Matoke-Muhia, D. et al. (2016) ‘Decline in frequency of the 2La chromosomal inversion in Anopheles gambiae (s.s.) in Western Kenya: Correlation with increase in ownership of insecticide-treated bed nets’, Parasites and Vectors, 9(1). Available at: 10.1186/s13071-016-1621-3.

Miles, A. et al. (2024) ‘malariagen/malariagen-data-python’: Zenodo. Available at: 10.5281/zenodo.13899118.

Mnzava, A.P. et al. (2015) ‘Implementation of the global plan for insecticide resistance management in malaria vectors: progress, challenges and the way forward’, Malaria Journal, 14(1), pp. 1–9. Available at: 10.1186/s12936-015-0693-4.

Morales-Pérezid, A. et al. (2020) ‘Utility of entomological indices for predicting transmission of dengue virus: Secondary analysis of data from the camino verde trial in Mexico and Nicaragua’, PLoS Neglected Tropical Diseases, 14(10), pp. 1–19. Available at: 10.1371/journal.pntd.0008768.

Neafsey, D.E., Taylor, A.R. and MacInnis, B.L. (2021) ‘Advances and opportunities in malaria population genomics’, Nature Reviews Genetics. Nature Research, pp. 502–517. Available at: 10.1038/s41576-021-00349-5.

Nei, M., Maruyama, T. and Chakraborty, R. (1975) ‘THE BOTTLENECK EFFECT AND GENETIC VARIABILITY IN POPULATIONS’, Evolution, 29(1), pp. 1–10. Available at: 10.1111/j.1558-5646.1975.tb00807.x.

Njoroge, H. et al. (2022) ‘Identification of a rapidly-spreading triple mutant for high-level metabolic insecticide resistance in Anopheles gambiae provides a real-time molecular diagnostic for anti-malarial intervention deployment’, Molecular Ecology [Preprint]. Available at: https://doi.org/10.1111/mec.16591.

O’Loughlin, S.M. et al. (2016) ‘Genomic signatures of population decline in the malaria mosquito Anopheles gambiae’, Malaria Journal, 15(1). Available at: 10.1186/s12936-016-1214-9.

Omondi, S. et al. (2023) ‘Late morning biting behaviour of Anopheles funestus is a risk factor for transmission in schools in Siaya, western Kenya’, Malaria Journal, 22(1). Available at: 10.1186/s12936-023-04806-w.

Praulins, G. et al. (2024) ‘Unpacking WHO and CDC Bottle Bioassay Methods: A Comprehensive Literature Review and Protocol Analysis Revealing Key Outcome Predictors’, Gates Open Research, 8. Available at: 10.12688/gatesopenres.15433.1.

Prussing, C. et al. (2018) ‘Decreasing proportion of Anopheles darlingi biting outdoors between long-lasting insecticidal net distributions in peri-Iquitos, Amazonian Peru’, Malaria Journal, 17(1). Available at: 10.1186/s12936-018-2234-4.

R Core Team (2019) ‘R: A Language and Environment for Statistical Computing’. Vienna, Austria: R foundation for statistical programming.

Ranson, H. et al. (2000) ‘Identification of a point mutation in the voltage-gated sodium channel gene of Kenyan Anopheles gambiae associated with resistance to DDT and pyrethroids’, Insect Mol Biol, 9. Available at: 10.1046/j.1365-2583.2000.00209.x.

Real, M.D., Ferre, J. and Chapa, F.J. (1991) ‘UDP-Glucosyltransferase Activity toward Exogenous Substrates in Drosophila melanogaster’, Analytical Biochemistry, (194), pp. 349–352.

Reidenbach, K.R. et al. (2014) ‘Cuticular differences associated with aridity acclimation in African malaria vectors carrying alternative arrangements of inversion 2La’, Parasites and Vectors, 7(1). Available at: 10.1186/1756-3305-7-176.

Rocca, K.A. et al. (2009) ‘2La chromosomal inversion enhances thermal tolerance of Anopheles gambiae larvae’, Malaria Journal, 8(1). Available at: 10.1186/1475-2875-8-147.

Rohani, A. et al. (2016) ‘Comparative Human Landing Catch and CDC Light Trap in Mosquito Sampling in Knowlesi Malaria Endemic Areas in Peninsula Malaysia’, Adv. Entomol., 4(January), pp. 1–10. Available at: 10.4236/ae.2016.41001.

Saarinen, E. V., Austin, J.D. and Daniels, J.C. (2010) ‘Genetic estimates of contemporary effective population size in an endangered butterfly indicate a possible role for genetic compensation’, Evolutionary Applications, 3(1), pp. 28–39. Available at: 10.1111/j.1752-4571.2009.00096.x.

Saarman, N.P. et al. (2017) ‘Effective population sizes of a major vector of human diseases, Aedes aegypti’, Evolutionary Applications, 10(10), pp. 1031–1039. Available at: 10.1111/eva.12508.

Schmidt, T.L., Endersby-Harshman, N.M. and Hoffmann, A.A. (2021) ‘Improving mosquito control strategies with population genomics’, Trends in Parasitology. Elsevier Ltd, pp. 907–921. Available at: 10.1016/j.pt.2021.05.002.

Staedke, S.G. et al. (2019) ‘Effect of long-lasting insecticidal nets with and without piperonyl butoxide on malaria indicators in Uganda (LLINEUP): a pragmatic, cluster-randomised trial embedded in a national LLIN distribution campaign’, Trials, 20(321). Available at: 10.1186/s13063-019-3382-8.

Staedke, S.G. et al. (2020) ‘Effect of long-lasting insecticidal nets with and without piperonyl butoxide on malaria indicators in Uganda (LLINEUP): a pragmatic, cluster-randomised trial embedded in a national LLIN distribution campaign’, The Lancet, 395(10232), pp. 1292–1303. Available at: 10.1016/S0140-6736(20)30214-2.

Strimmer, K. (2008) ‘fdrtool: A versatile R package for estimating local and tail area-based false discovery rates’, Bioinformatics, 24(12), pp. 1461–1462. Available at: 10.1093/bioinformatics/btn209.

Swai, J.K. et al. (2023) ‘CDC light traps underestimate the protective efficacy of an indoor spatial repellent against bites from wild Anopheles arabiensis mosquitoes in Tanzania’, Malaria Journal, 22(1). Available at: 10.1186/s12936-023-04568-5.

The Anopheles gambiae 1000 Genomes Consortium (2017) ‘Genetic diversity of the African malaria vector Anopheles gambiae’, Nature Research Letter [Preprint]. Available at: 10.1038/nature24995.

Topham, M.K. (2006) ‘Signaling roles of diacylglycerol kinases’, Journal of Cellular Biochemistry, pp. 474– 484. Available at: 10.1002/jcb.20704.

Traore, M.M. et al. (2020) ‘Large-scale field trial of attractive toxic sugar baits (ATSB) for the control of malaria vector mosquitoes in Mali, West Africa’, Malaria journal, 19(1), pp. 1–16.

Vontas, J. et al. (2018) ‘Rapid selection of a pyrethroid metabolic enzyme CYP9K1 by operational malaria control activities’, Proceedings of the National Academy of Sciences of the United States of America, 115(18), pp. 4619–4624. Available at: 10.1073/pnas.1719663115.

Weetman, D. et al. (2018) ‘Candidate-gene based GWAS identifies reproducible DNA markers for metabolic pyrethroid resistance from standing genetic variation in East African Anopheles gambiae’, Scientific Reports, 8(1), pp. 1–12. Available at: 10.1038/s41598-018-21265-5.

Weetman, D. and Donnelly, M.J. (2015) ‘Evolution of insecticide resistance diagnostics in malaria vectors’, Trans. R. Soc. Trop. Med. Hyg., 109(5), pp. 291–293. Available at: 10.1093/trstmh/trv017.

WHO (2024) World Malaria Report 2024. Geneva: World Health Organization.

Wilson, A.L. et al. (2020) ‘The importance of vector control for the control and elimination of vector-borne diseases’, PLoS Neglected Tropical Diseases. Public Library of Science, pp. 1–31. Available at: 10.1371/journal.pntd.0007831.

