## Supplementary Fig 1 for "Genetic Surveillance Reveals Differential Evolutionary Dynamic of *Anopheles gambiae* Under Contrasting Insecticidal Tools used in Malaria control"

**Supplementary figure 1**

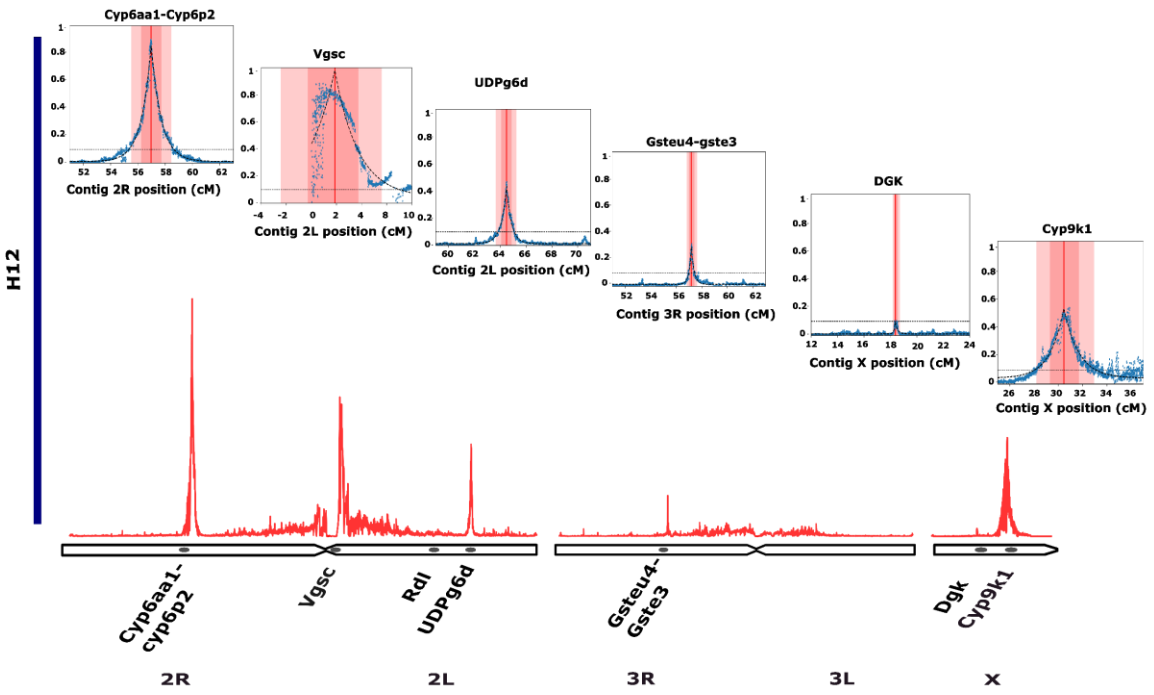


***Supplementary Fig.1 Genomic regions under selection in Anopheles gambiae populations in Uganda.*** *H12 analysis identified genomic regions under selection, and an exponential modelling approach was used to determine peak centres, guiding the selection of regions for haplotype cluster analysis. Apart from the sweep in the Gstu4-Gste3 region, the frequencies of dominant haplotypes in other regions changed significantly over the course of the trial and were included in the haplotype clustering analysis*
