## Supplementary Fig 2 for "Genetic Surveillance Reveals Differential Evolutionary Dynamic of *Anopheles gambiae* Under Contrasting Insecticidal Tools used in Malaria control"

**Supplementary figure 2**

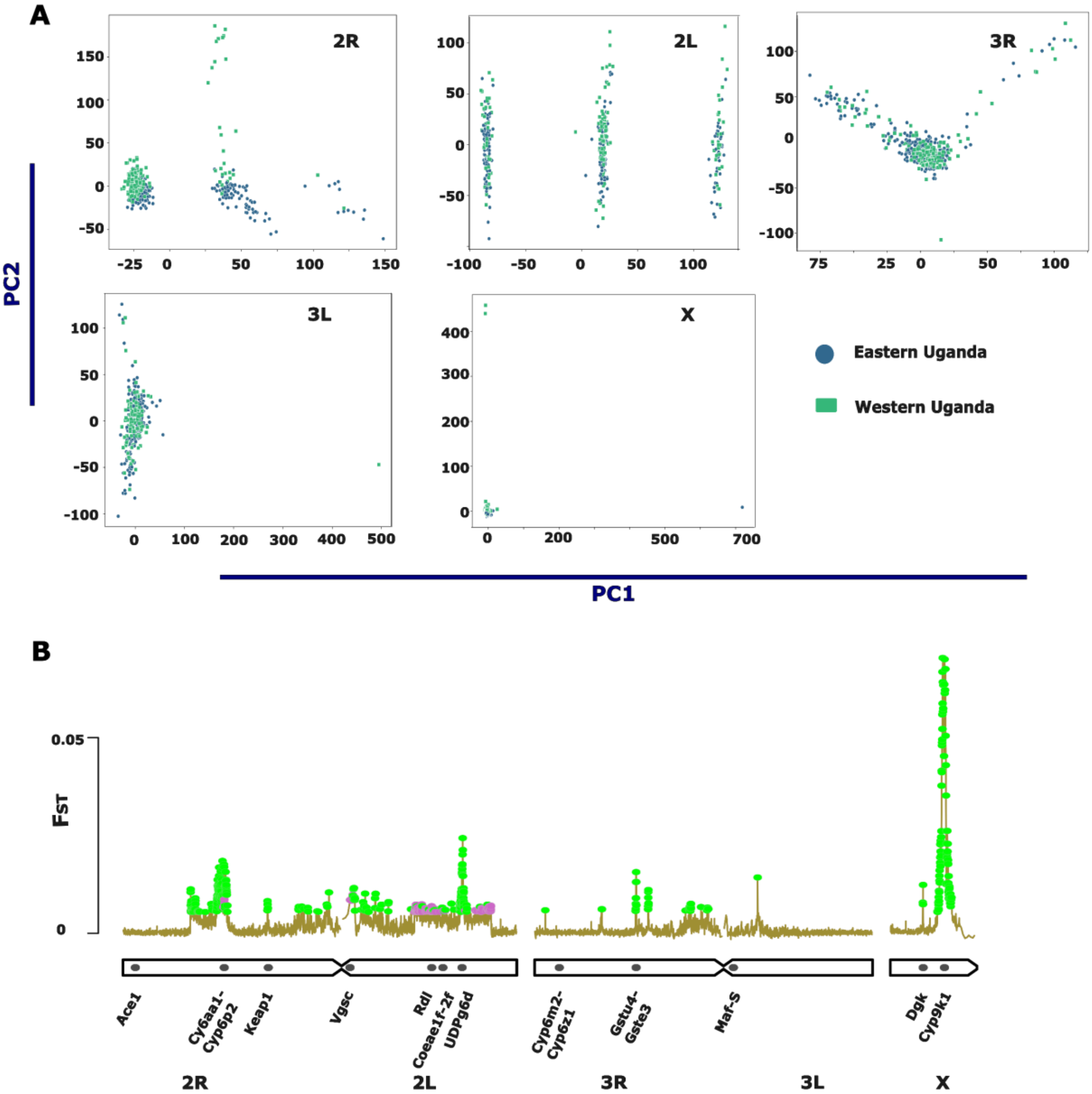


***Supplementary Fig.2 Population structure of Anopheles gambiae population in Uganda.***

***A:*** *Principal component analysis of An. gambiae population structure before net distribution based on chromosomes 2R, 2L, 3R, 3L and X did not reveal population structure between mosquitoes collected in Eastern vs Western Uganda but structured in chromosome 2R and 2L based on chromosomal inversions (2Rb and 2La respectively).* ***B:*** *Genome wide windowed* $F_{ST}$ *analysis revealed population differentiation between the two populations mostly in loci associated with insecticide resistance. On the* $F_{ST}$ *plot, colored dots mark windows surpassing the permutation-based significance threshold (green: significant; purple: marginal).*
