## Supplementary Fig 3 for "Genetic Surveillance Reveals Differential Evolutionary Dynamic of *Anopheles gambiae* Under Contrasting Insecticidal Tools used in Malaria control"

**Supplementary figure 3**

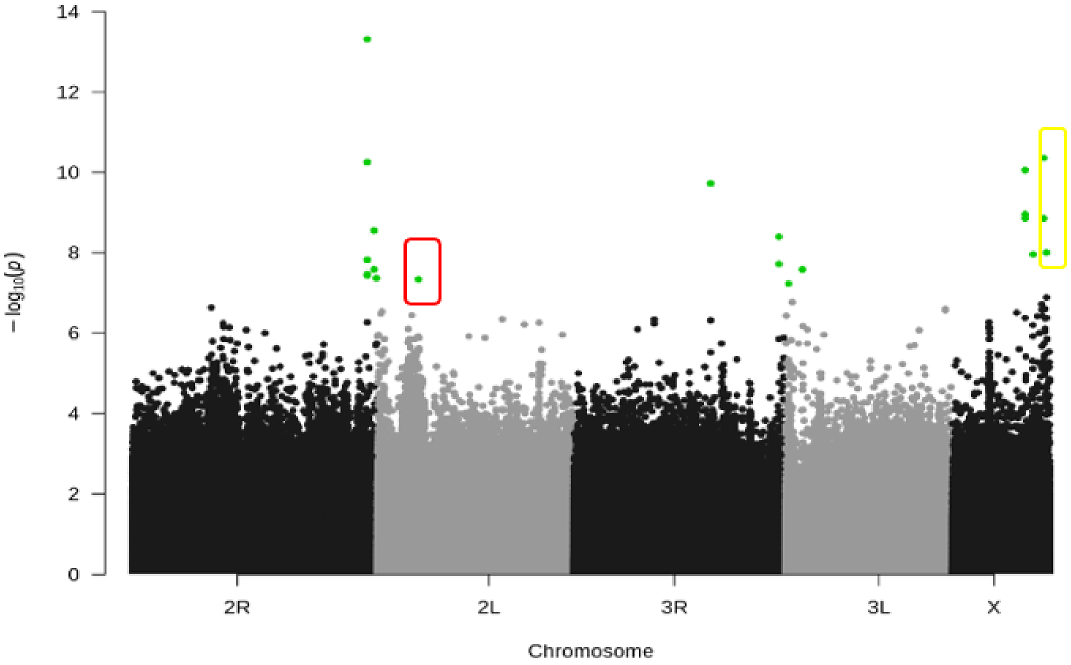


***Supplementary Fig.3*** *A genome wide analysis of temporal variation in SNP frequency in Anopheles gambiae population driven by either pyrethroid exposure in standard pyrethroid only nets or Pyrethroid-PBO nets distributed in Uganda. The highlighted P values (green) are SNPs that changed significantly during the intervention (FDR <0.05). One SNP in a red rectangle (2L:10334049, A>C, V390G) in AGAP005127 (RNA-binding protein 15) was non-synonymous, while three were intronic SNPs (in yellow rectangle) within CYP4G16 (X:22941281 and X:22941291) and Fatty acyl-CoA reductase (X:23560372). The remaining SNPs were intergenic.*
