## Supplementary Fig 4 for "Genetic Surveillance Reveals Differential Evolutionary Dynamic of *Anopheles gambiae* Under Contrasting Insecticidal Tools used in Malaria control"

**Supplementary figure 4**

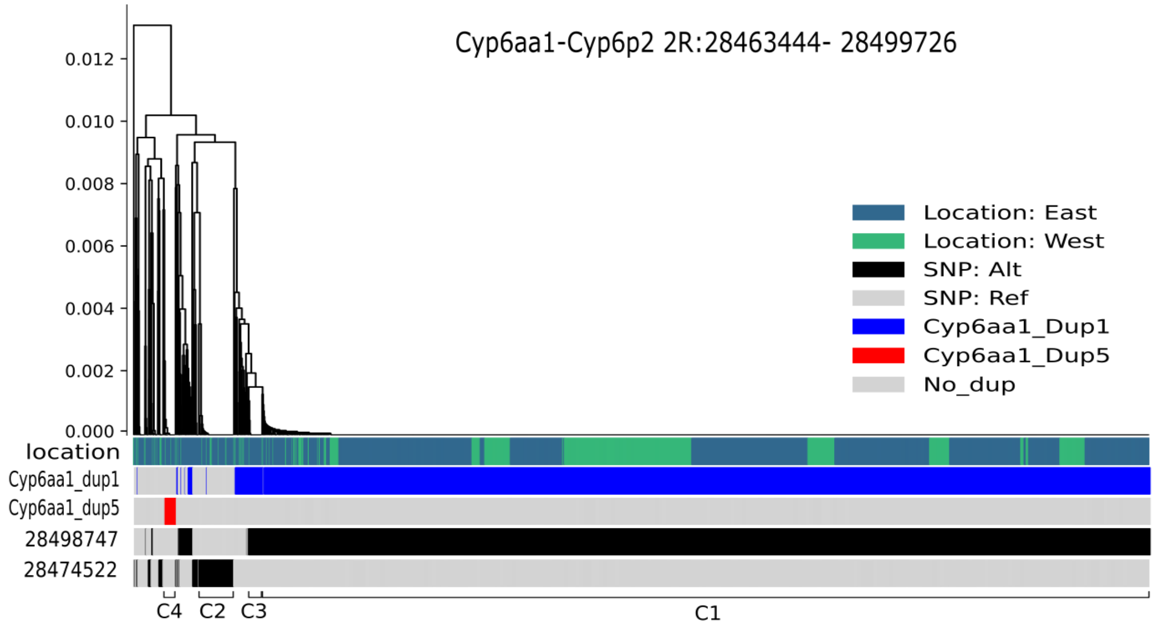


***Supplementary Fig.4 Haplotype dendrogram of the Cyp6aa1-Cyp6p2 region showing the main haplotype is associated with Cyp6aa1 duplication.*** *Each dendrogram leaf represents a haplotype. Coloured bars beneath each dendrogram show for each haplotype it's geographical origin (green = Western Uganda, navy blue = Eastern Uganda) and the presence (blue / red) or absence (light grey) of known CNV alleles on that haplotype. SNPs tagging the haplotype clusters are represented in black colour while reference allele is light grey.*
