## Supplementary Table 1 for "Genetic Surveillance Reveals Differential Evolutionary Dynamic of *Anopheles gambiae* Under Contrasting Insecticidal Tools used in Malaria control"

***Supplementary Table 1. The number of samples used in the genomic analysis.*** *Because of the low number of intermediate collections across the 4 cohorts, only baseline and round 5 (25 months post intervention) were included in the analysis.*

| Location | Trial arm | Baseline | Round 2 | Round 3 | Round 4 | Round 5 | Total |
|---|---|---|---|---|---|---|---|
| **East** | Nonpbo | **132** | 47 | 148 | 65 | **192** | 584 |
| | Pbo | **112** | 0 | 10 | 1 | **39** | 162 |
| **West** | Nonpbo | **75** | 3 | 6 | 61 | **30** | 175 |
| | Pbo | **73** | 1 | 4 | 0 | **14** | 92 |
