## Supplementary Table 2 for "Genetic Surveillance Reveals Differential Evolutionary Dynamic of *Anopheles gambiae* Under Contrasting Insecticidal Tools used in Malaria control"

***Anopheles density ratios over time after LLIN distribution***

| ROUND | DENSITY RATIO PBO BEDNETS | DENSITY RATIO STANDARD BEDNETS | EFFECT SIZE* | P VALUES* |
|---|---|---|---|---|
| BASELINE | Reference | Ref. | Ref. | Ref. |
| 6 MONTHS | 0.2129103 | 0.8683859 | -0.6221 | **0.000326** |
| 12 MONTHS | 0.3625230 | 1.0792079 | -0.4677 | **0.000539** |
| 18 MONTHS | 0.4802311 | 1.3610011 | -0.2093 | 0.27 |
| 25 MONTHS | 0.6924962 | 1.7537129 | -0.1156 | 0.36 |

**The effect size and p value were calculated using a generalised linear mixed model in which the net type and round were fixed effects while household id was fitted as a random effect.*
