## Supplementary Table 3 for "Genetic Surveillance Reveals Differential Evolutionary Dynamic of *Anopheles gambiae* Under Contrasting Insecticidal Tools used in Malaria control"

***Population genetics analysis to infer changes in An. gambiae population size after Bed net distribution during the LLINEUP trial in Uganda***

| Region | Cohort | LD[a] | Ne[b] | Nucleotide diversity (π)[c] |
|---|---|---|---|---|
| All | All baseline | 0.0007[0.00005, 0.015] | Inf [3636, Inf] | 0.0214 [0.0201, 0.0226] |
| | All post | 0.0007[0.00005, 0.015] | Inf [4036, Inf] | 0.0211 [0.0199, 0.0224] |
| | PBO baseline | 0.007[0.0003, 0.08] | 3348 [1273, Inf] | 0.0212 [0.0199, 0.0224] |
| | PBO post | 0.007[0.0003, 0.08] | Inf [1554, Inf] | 0.0211 [0.0199, 0.0224] |
| | Standard baseline | 0.001[0.0001, 0.02] | 7210 [3169, Inf] | 0.0215 [0.0202, 0.0227] |
| | Standard post | 0.001[0.0001, 0.02] | Inf [3478, Inf] | 0.0211 [0.0199, 0.0224] |
| West | PBO baseline | 0.051[1.7e-32, 0.412] | 1008 [229, Inf] | 0.0212 [0.020, 0.0225] |
| | PBO post | 0.051[2.3e-32, 0.412] | Inf [244, Inf] | 0.0210 [0.0197, 0.0223] |
| | Standard baseline | 0.017[0.00009, 0.209] | 3408 [708, Inf] | 0.0211 [0.0199, 0.0224] |
| | Standard post | 0.017[0.00009, 0.210] | Inf [702, inf] | 0.0211 [0.0199, 0.0224] |
| East | PBO baseline | 0.011[0.00008, 0.147] | 949 [853, Inf] | 0.0212 [0.0199, 0.0224] |
| | PBO post | 0.011[0.00008, 0.149] | 12876 [667, Inf] | 0.0211 [0.0199, 0.0224] |
| | Standard baseline | 0.002[0.00004, 0.053] | Inf [2429, Inf] | 0.0211 [0.0199, 0.0224] |
| | Standard post | 0.002[0.00004, 0.053] | Inf [2176, Inf] | 0.0211 [0.0199, 0.0224] |

*[a] LD is the median of R squares calculated from squared correlation coefficient ($r^2$) for SNPs pairs within chromosomes 2 vs 3. In brackets are the 5 and 95% percentiles. [b]Ne is an estimate from LD and in brackets are 95% CI estimated from 1000 replicates. [c]π is the nucleotide diversity estimate and in brackets 95% CI.*
