## Supplementary Table 4 for "Genetic Surveillance Reveals Differential Evolutionary Dynamic of *Anopheles gambiae* Under Contrasting Insecticidal Tools used in Malaria control"

***Haplotype Cluster Analysis Highlights Region- and Net-Specific Genetic Responses in Anopheles gambiae to Bed Net Interventions.***

| Region | Hypothesis | C1 | | C2 | | C3 | | C4 | |
|---|---|---|---|---|---|---|---|---|---|
| | | p | coef | p | coef | p | coef | p | coef |
| ***Cyp6aa1-Cyp6p2*** | $H_3$[a] | **0.02** | 0.4 | **0.002 [0.02]** | -1.02 | 0.28 | 0.58 | 0.98 | -0.013 |
| | $H_4$[b]-PBO | 0.064 | 0.7 | **0.0022 [0.02]** | -48 | **0.03** | 2.3 | 0.8 | 0.32 |
| | $H_4$[b]-standard | 0.12 | 0.31 | **0.03** | -0.78 | 0.93 | -0.055 | 0.94 | -0.043 |
| ***Vgsc*** | $H_3$ | 0.3 | -0.12 | 0.38 | -0.13 | 0.08 | 0.68 | **0.012** | 1.3 |
| | $H_3$[c]-East | 0.37 | -0.12 | 0.51 | 0.09 | 0.08 | 0.9 | 0.36 | 1.1 |
| | $H_3$[c]- West | 0.79 | -0.07 | **0.007** | -0.8 | 0.69 | 0.22 | **0.02** | 1.3 |
| | $H_4$- PBO | 0.063 | -0.42 | 0.91 | 0.026 | 0.36 | 0.6 | 0.13 | 1.1 |
| | $H_4$-standard | 0.92 | 0.015 | 0.1 | -0.29 | 0.11 | 0.77 | **0.029** | 1.8 |
| ***2L-34mb*** | $H_3$ | 0.35 | 0.16 | 0.65 | 0.12 | 0.074 | -0.86 | 0.47 | 0.51 |
| | $H_4$-PBO | **0.0003 [0.003]** | 0.9 | 0.54 | -0.31 | 0.89 | 0.15 | 0.34 | -19 |
| | $H_4$-standard | 0.7 | -0.08 | 0.49 | 0.21 | **0.039** | -1.1 | 0.12 | 1.4 |
| ***Dgk*** | $H_3$ | **0.0004 [0.003]** | 0.87 | 0.83 | -0.05 | 0.053 | -0.96 | 0.69 | 0.28 |
| | $H_4$-PBO | 0.42 | 0.37 | 0.94 | 0.031 | 0.75 | -0.25 | 0.32 | -20 |
| | $H_4$-standard | **2.7e-05 [0.0005]** | 1.1 | 0.69 | -0.098 | **0.024** | -1.4 | 0.63 | 0.4 |
| ***Cyp9k1*** | $H_3$ | 0.22 | 0.18 | 0.91 | -0.03 | 0.95 | -0.02 | 0.11 | -0.84 |
| | $H_4$-PBO | 0.29 | -0.32 | 0.4 | 0.43 | 0.53 | -0.38 | 0.88 | 0.13 |
| | $H_4$-standard | **0.0089 [0.04]** | 0.44 | 0.34 | -0.24 | 0.49 | 0.34 | 0.063 | 1.4 |

*In bold are raw p values of haplotypes that changed significantly (<0.05) but after FDR correction only those with the square brackets were below the set significance threshold (5%).*

[a] *model used to test $H_3$: variant ~ Round (1 vs 5) + location (east & west) + llin (PBO & standard) + 1|health subdistrict*

[b] *model used to test $H_4^b$: variant ~ Round (1 vs 5) + location (east & west) + 1|health subdistrict (Data split by net type)*

[c] *model used to test $H_4^c$: variant ~ Round (1 vs 5) + llin (PBO & standard) + 1|health subdistrict (Data split by location)*
