## Supplementary Table 5 for "Genetic Surveillance Reveals Differential Evolutionary Dynamic of *Anopheles gambiae* Under Contrasting Insecticidal Tools used in Malaria control"

***Top Single nucleotide polymorphism (SNPs) associated with main haplotypes in genomic regions where significant changes occurred during the LLINEUP bed net trial in Uganda***

| Region | Haps_C1 | Haps_C2 |
|---|---|---|
| **2R:28463444–28499726** | 2R:28500104<br>2R:28499987<br>2R:28499354<br>2R:28498747<br>2R: 28497958 | 2R:28476019<br>2R:28475974<br>2R:28474555<br>2R:28474522<br>2R:28507610 |
| **2L:2791320–2893275** | 2L:2883338<br>2L:2569930<br>2L:2300012<br>2L:2300046 | 2L:2409947<br>2L:2380726<br>2L:2383037<br>2L:2321129 |
| **2L:34081017–34101131** | 2L:34076000<br>2L: 34073954<br>2L:34093806<br>2L:34100535<br>2L:34103278 | 2L:34100533<br>2L:34098712<br>2L:34097024<br>2L:34098608<br>2L:34096945 |
| **X:9179019–9185374** | X:9179818<br>X:9183213<br>X:9183402<br>X:9190710 | X:9185342<br>X:9183189<br>X:9181789<br>X:9181138 |
| **X:15216225–15271654** | X:15239300<br>X:15238600<br>X:15237378<br>X:15236958<br>X:15236122 | X:15247409<br>X:15247243<br>X:15242464<br>X:15238589<br>15206248 |
