## Supplementary |Table 6 for "Genetic Surveillance Reveals Differential Evolutionary Dynamic of *Anopheles gambiae* Under Contrasting Insecticidal Tools used in Malaria control"

**Change in Knockdown resistance (KDR) haplotypes during the LLINEUP bednet intervention in Uganda.**

| *Kdr haplotypes* | *H3* | | *H3_east* | | *H3_west* | | *H4 PBO* | | *H4 non-PBO* | |
|---|---|---|---|---|---|---|---|---|---|---|
| | p | coef | p | coef | p | coef | p | coef | p | coef |
| *S1* | 0.17 | -0.2 | 0.47 | -0.1 | 0.2 | -0.3 | **0.04 [0.3]** | -0.5 | 0.86 | -0.03 |
| *S3* | 0.28 | -0.2 | 0.9 | 0.01 | 0.05 [0.3] | -0.6 | 0.65 | -0.1 | 0.15 | -0.3 |
| *S4* | - | - | - | - | - | - | - | - | 1 | -6.3 |
| *S* | 0.77 | 0.2 | 0.7 | -0.4 | 0.33 | 1 | 0.12 | 1.3 | 0.33 | -1.3 |
| *F3* | 0.22 | 1.3 | - | - | 0.22 | 1.3 | **0.007 [0.049]** | 33 | - | - |
| *F5* | **0.006 [0.04]** | 1.8 | 0.09 | 21 | **0.01 [0.08]** | 1.6 | **0.04 [0.3]** | 1.8 | **0.02 [0.1]** | 3 |
| *F* | 0.33 | 0.6 | 0.43 | -1.8 | 0.28 | 0.7 | 0.5 | 0.6 | 0.62 | 0.5 |
| *WT* | 0.4 | -0.5 | 0.18 | -1 | 0.4 | 0.8 | 0.34 | -0.7 | 0.64 | -0.4 |

*We tested two hypotheses: H3 -The pyrethroid in both net types will drive similar changes in KDR haplotypes frequencies. This was tested in both regions combined or separately. H4- Net specific changes in KDR haplotype frequencies will be observed. The p values in bold are where the haplotype change was significant (in square brackets) after correction for multiple testing (Bonferroni = 5%)
